# Optimization of Golden Gate assembly through application of ligation sequence-dependent fidelity and bias profiling

**DOI:** 10.1101/322297

**Authors:** Potapov Vladimir, Jennifer L. Ong, Rebecca B. Kucera, Bradley W. Langhorst, Katharina Bilotti, John M. Pryor, Eric J. Cantor, Barry Canton, Thomas F. Knight, Thomas C. Evans, Gregory J. S. Lohman

## Abstract

Modern synthetic biology depends on the manufacture of large DNA constructs from libraries of genes, regulatory elements or other genetic parts. Type IIS restriction enzyme-dependent DNA assembly methods (e.g., Golden Gate) enable rapid one-pot, ordered, multi-fragment DNA assembly, facilitating the generation of high-complexity constructs. The order of assembly of genetic parts is determined by the ligation of flanking Watson-Crick base-paired overhangs. The ligation of mismatched overhangs leads to erroneous assembly, and the need to avoid such pairings has typically been accomplished by using small sets of empirically vetted junction pairs, limiting the number of parts that can be joined in a single reaction. Here, we report the use of a comprehensive method for profiling end-joining ligation fidelity and bias to predict highly accurate sets of connections for ligation-based DNA assembly methods. This data set allows quantification of sequence-dependent ligation efficiency and identification of mismatch-prone pairings. The ligation profile accurately predicted junction fidelity in ten-fragment Golden Gate assembly reactions, and enabled efficient assembly of a *lac* cassette from up to 24-fragments in a single reaction. Application of the ligation fidelity profile to inform choice of junctions thus enables highly flexible assembly design, with >20 fragments in a single reaction.

## INTRODUCTION

The one-pot assembly of large DNA structures from smaller component parts is a key technology in modern synthetic biology, with common *in vitro* methods dependent on high-fidelity ligation steps to produce the desired constructs. Accurate ligation is essential to Gibson assembly and related technologies, which require the joining of fragments at long overlapping regions (1,2). Low fidelity ligation could lead to the inclusion of mismatches, deletions or insertions in final assemblies, lowering overall yield of full length construct and increasing the chances of a given colony having one or more mistakes. In the restriction enzyme-dependent assembly methods such as BioBricks™ and Golden Gate cloning, the assembly of large constructs is achieved by the sequential or parallel joining of multiple DNA fragments linked by short overhangs (3–7). To produce the desired final assembly, fragments must be joined only by Watson-Crick overhang pairs; if mismatched overhangs ligate, incorrect assemblies will result with large insertions or deletions of entire fragments, or result in one or more fragments being inserted in the incorrect orientation.

T4 DNA ligase is commonly employed in Type IIS restriction enzyme-dependent DNA assembly methods, which allow the joining of multiple fragments in one pot, due to its high efficiency in end-joining reactions. However, this enzyme is well known to join mismatches, gaps and other imperfect structures with varying levels of efficiency (8–12). In order to assure high-fidelity assembly in Golden Gate and derived methods, several rules of thumb have been adopted to minimize the risk of ligating imperfectly base-paired partners during an assembly reaction. In addition to the obvious need to not use any overhangs more than once within an assembly, and to avoid palindromic overhangs to prevent self-ligation of fragments, it is typically advised to have at least a two base-pair difference between overhangs, to avoid overhangs with three identical base-pairs in a row, and to ensure all overhangs have similar GC content in a given assembly (4–6). Following these rules restricts the user to a limited number of four-base overhangs, which is particularly constraining when sequences may not be chosen arbitrarily (e.g., when assemblies must break within coding sequences). In addition, several systematic Golden Gate based assembly methods (e.g., MoClo, Golden Braid, Mobius Assembly, MIDAS), have further restricted the number of overhangs to standardized, reliable sets in an effort to improve efficiency and fidelity (13–20). While very large DNA constructs can be produced from successive hierarchical assembly rounds in these methods, the number of fragments that can be assembled in a single pot is limited by the number of allowable overhang pairs, typically limiting the user to six to eight fragments at a time. Both the rules of thumb and the standardized sets were developed based on empirical but non-comprehensive studies, and the possibility exists to greatly expand the power of these methods through the Identification of additional high-fidelity overhang sets, especially larger sets, with limited expected mismatch ligation between any two overhangs.

Here, we report the application of a single-molecule, next-generation sequencing assay to probe the fidelity of DNA ligase end-joining in the context of four-base overhangs (Figure 1). Applying our previously reported sequencing method for profiling ligase end-joining fidelity (21), we have quantified the ligation efficiency of all four-base Watson-Crick pairs and the prevalence of all possible mismatched overhang combinations by T4 DNA ligase at 25 and 37°C with typical buffer conditions lacking molecular crowding agents. We have further applied this data to predict the accuracy of a ten-fragment Golden Gate assembly, demonstrating the ability to predict overall assembly fidelity, specific assembly errors, and ligation pairs that exhibit relatively low ligation efficiency despite perfectly complementary overhangs. Finally, we apply the ligation data to guide the choice of ligation junctions in the design of 12- and 24-fragment assemblies of a *lac* cassette. Efficient assembly of the correct gene construct was observed when predicted high-fidelity junction sets are used, and a designed deletion-prone, 12-fragment assembly likewise closely matched predictions. The current data set can thus be used to enumerate sets of “orthogonal” junctions, with little to no predicted mismatch ligation amongst any members of the set, allowing >20 fragment, one-pot assemblies with much greater flexibility in choosing junctions than allowed by traditional rules of thumb. We further report optimal high-fidelity junction sets enumerated from the data, predicted to allow high-fidelity assembly when sequences may be chosen arbitrarily.

**Figure 1.**
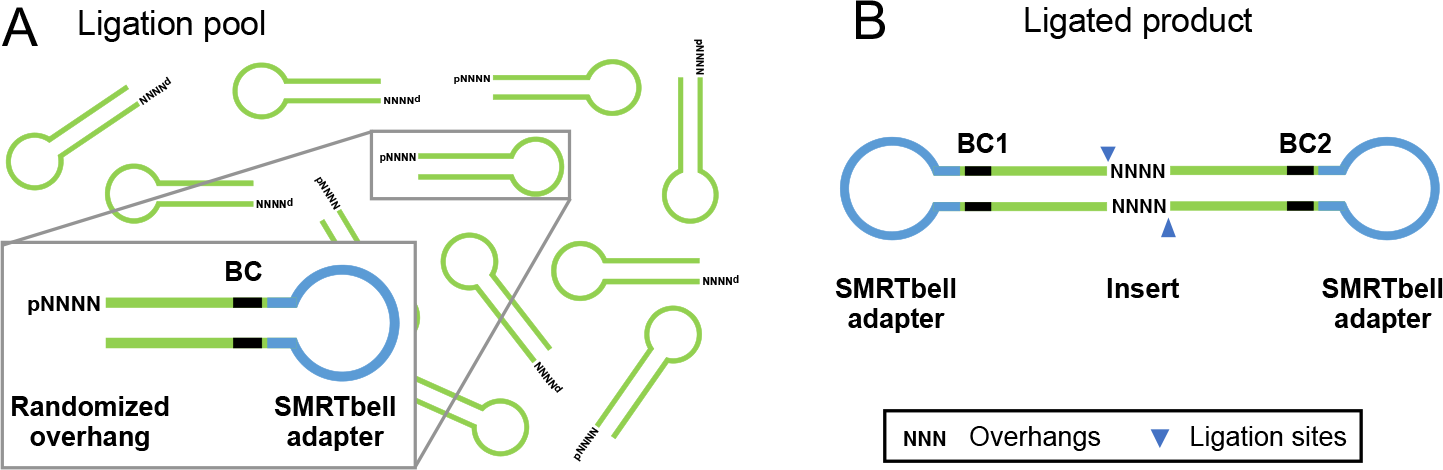
Schematic of multiplexed ligation fidelity and bias profiling assay. (A) Libraries containing randomized four-base overhangs were synthesized and ligated with T4 DNA ligase under various conditions. The hairpin substrates contain the Pacific-Biosciences SMRTbell adapter sequence, an internal 6-base random barcode used to confirm strand identity and monitor the substrate sequence bias derived from oligonucleotide synthesis, and randomized four-base overhangs. (B) Ligated substrates form circular molecules, in which a double-stranded insert DNA is capped with SMRTbell adapters. These products were sequenced utilizing Pacific Biosciences SMRT sequencing, which produced long rolling-circle sequencing reads. Consensus sequences were built for the top and bottom strands independently, allowing extraction of the overhang identity and barcode sequence.

## MATERIAL AND METHODS

All enzymes and buffers were obtained from New England Biolabs (NEB, Ipswich, MA) unless otherwise noted. T4 DNA ligase reaction buffer (1X) is: 50 mM Tris-HCl (pH 7.5), 10 mM MgCl_2_, 1 mM ATP, 10 mM DTT. NEBuffer 2 (1X) is: 10 mM Tris-HCl (pH 7.9), 50 mM NaCl, 10 mM MgCl_2_, 1 mM DTT. CutSmart Buffer (1X) is: 20 mM Tris-acetate (pH 7.9), 50 mM Potassium Acetate, 10 mM Magnesium Acetate, 100 μg/ml BSA. Thermopol buffer is: 20 mM Tris-HCl (pH 8.8), 10 mM (NH_4_)_2_SO_4_, 10 mM KCl, 2 mM MgSO_4_, 0.1% Triton^®^-X-100. Standard Taq polymerase buffer is: 10 mM Tris-HCl (pH 8.3), 50 mM KCl, 1.5 mM MgCl_2_. SOC outgrowth medium and competent *E. coli* strain T7 Express were from New England Biolabs. The T7 express cell line lacks a functional *lacZ* gene, full genotype: *fhuA2 lacZ::T7 gene1 [lon] ompT gal sulA11 R(mcr-73::miniTn10--*Tet^S^*)2 [dcm] R(zgb-210::Tn10--*Tet^S^*) endA1 Δ(mcrC-mrr)114::IS10*. All column cleanup of oligonucleotides and ligated libraries was performed using Monarch^®^ PCR & DNA Cleanup Kit columns (NEB), following the published Oligonucleotide Cleanup Protocol (https://www.neb.com/protocols/2017/04/25/oligonucleotide-cleanup-using-monarch-pcr-dna-cleanup-kit-5-g-protocol-neb-t1030). Oligonucleotide purity and sizing was performed using an Agilent Bioanalyzer 2100, using a DNA 1000 assay, following the standard protocols. Synthetic oligonucleotides were obtained from Integrated DNA Technologies as lyophilized solid (Coralville, IA). Fragments for Golden Gate assembly assays were obtained from GenScript (Piscataway, NJ), as precloned inserts flanked by BsaI cut sites in a pUC57-mini plasmid with the native BsaI site in the *amp^R^* gene removed through silent mutagenesis.

### Preparation and Pacific Biosciences SMRT sequencing of ligation fidelity libraries

The substrate for the four-base overhang ligation fidelity assay was produced using the protocol previously published for three-base overhangs, with the following modifications (21). Initial PAGE-purified substrate precursor oligonucleotide (IDT) contained a 5′-terminal region, a randomized four-base region, a BsaI binding site, a constant region, an internal 6-base randomized region as a control for synthesis bias, and a region corresponding to the SMRT-bell sequencing adapter for Pacific Biosciences SMRT sequencing (Supplementary Data, Table S1). The oligonucleotide was designed with a short (7-base) complementary region such that they form a primer-template junction hairpin structure (Figure 1). The precursor oligonucleotide was extended as per the published method (21). The extended DNA was purified (Monarch^®^ PCR & DNA Cleanup Kit), and the concentration of the purified DNA (typically 25 – 30 μM) was determined using an Agilent Bioanalyzer 2100, DNA 1000 kit.

The extended DNA was cut using BsaI to generate a four-base overhang. Typically, 1 μL of DNA from the extension reaction was combined with 900 U of BsaI in a 100 μL total volume of NEB CutSmart buffer and incubated for 2 h at 37°C. Reactions were halted by addition of 1 μL Proteinase K followed by 20 min incubation at 37°C, then purified using the Monarch^®^ PCR & DNA Cleanup Kit (NEB). Final concentration and extent of cutting was determined by Agilent Bioanalyzer (DNA 1000) and confirmed to be >95% cut. Remaining uncut starting material (~5%) was not 5′ phosphorylated and thus should not interfere with subsequent cohesive-end joining reactions. For use in subsequent steps, DNA substrates were diluted to ~500 nM in 1X TE buffer, with precise concentration determined by Bioanalyzer. The final substrate sequence can be found in the Supplementary Data, Table S1.

In a typical ligation reaction, substrate (100 nM) was combined with 2.5 μL high concentration T4 DNA ligase (2000 U, 1.75 μM final concentration) in 1× T4 DNA ligase buffer in a 50 μL total reaction volume and incubated for 1 h or 18 h at 25°C or 37°C. Reactions were quenched with 2.5 μL 500 mM EDTA, and purified using the Monarch^®^ PCR & DNA Cleanup Kit, oligonucleotide cleanup protocol. Each ligation was performed in a minimum of duplicates, and the ligation yield was determined by Agilent Bioanalyzer (DNA 1000) with error reported as one standard deviation. The ligated library was treated with Exonuclease III (50U) and Exonuclease VII (5 U) in a 50 μL volume in 1X Standard Taq Polymerase buffer for a 1 h incubation at 37°C. The library was purified using a Monarch^®^ PCR & DNA Cleanup Kit, oligonucleotide cleanup protocol, including a second wash step, then quantified by Agilent Bioanalyzer (DNA 1000). Typical concentrations of final library were between 0.5 and 2 ng/μL. Sequencing and analysis of sequencing data were performed as previously described (21), with the scripts modified to use the expected insert sequence from the four-base overhang ligation reactions (Supplementary Data, Table S1).

### Preparation and SMRT sequencing analysis of ten-fragment Golden Gate assemblies

Randomized insert sequences were designed by combining all possible 4-base sequences *in sillico* in random order to generate 1024nt sequences. Three such sequences were combined, then divided into ten 300nt fragments (discarding the remaining 72nt fragment). Homopolymer regions of length greater than 4 and BsaI restriction sites were excluded from all sequences. The final ten fragments (Supplementary Data, Table S2) were obtained precloned into pUC57-mini plasmids, flanked with BsaI cut sites designed to allow arbitrary 4-base overhangs to be installed on the ends of each insert; the junction overhangs used in each assembly set are specified in Table 1.

**Table 1.** Ten-fragment Golden Gate assembly junction sequences.

| Junction | High-fidelity | Deletion-prone | Failure-prone | Low-fidelity |
| --- | --- | --- | --- | --- |
|  | set | set | set | set |
| $\frac{1}{1'}$ | $\frac{AAGG}{TTCC}$ | $\frac{\text{-----}}{\text{-----}}$ | $\frac{\text{-----}}{\text{-----}}$ | $\frac{GCCC}{CGGG}$ |
| $\frac{2}{2'}$ | $\frac{ACTC}{TGAG}$ | $\frac{\text{-----}}{\text{-----}}$ | $\frac{\text{-----}}{\text{-----}}$ | $\frac{GCCA}{CGGT}$ |
| $\frac{3}{3'}$ | $\frac{AGGA}{TCCT}$ | $\frac{\text{-----}}{\text{-----}}$ | $\frac{\text{-----}}{\text{-----}}$ | $\frac{ACCC}{TGGG}$ |
| $\frac{4}{4'}$ | $\frac{AGTC}{TCAG}$ | $\frac{\text{-----}}{\text{-----}}$ | $\frac{\text{-----}}{\text{-----}}$ | $\frac{AGCC}{TCGG}$ |
| $\frac{5}{5'}$ | $\frac{ATCA}{TAGT}$ | $\frac{\text{-----}}{\text{-----}}$ | $\frac{\text{-----}}{\text{-----}}$ | $\frac{CGCC}{GCGG}$ |
| $\frac{6}{6'}$ | $\frac{GCCG}{CGGC}$ | $\frac{\text{-----}}{\text{-----}}$ | $\frac{\text{-----}}{\text{-----}}$ | $\frac{AGCA}{TCCT}$ |
| $\frac{7}{7'}$ | $\frac{CTGA}{GACT}$ | $\frac{GCTG}{CGAC}$ | $\frac{TAAA}{ATTT}$ | $\frac{AGCG}{TCGC}$ |
| $\frac{8}{8'}$ | $\frac{GCGA}{CGCT}$ | $\frac{\text{-----}}{\text{-----}}$ | $\frac{\text{-----}}{\text{-----}}$ | $\frac{CGGC}{GCCG}$ |
| $\frac{9}{9'}$ | $\frac{GGAA}{CCTT}$ | $\frac{\text{-----}}{\text{-----}}$ | $\frac{\text{-----}}{\text{-----}}$ | $\frac{AGGC}{TCCG}$ |

<sup>1</sup> A notation of $\frac{\text{-----}}{\text{-----}}$ indicates the junction pair used is identical to the HF set.

Insert fragments were amplified using PCR primers designed to anneal to the plasmid region flanking the insert site (P1 GGGTTCCGCGCACATTTC; P2 TTTGCTGGCCTTTTGCTCACAT). PCR reactions included 100 pg/μL plasmid, 0.5 μM each primer, 2 U Q5 High Fidelity DNA polymerase, and 0.2 mM each dNTP in Q5 Reaction buffer in a 100 μL total reaction volume. Reactions were incubated 30 s at 98°C, 16 cycles of 5 s at 98°C, 10 s at 62°C, and 20 s at 72°C, with a final incubation of 5 min at 72°C, with the exception of insert D for the low-fidelity set, which was amplified by EpiMark Hot Start *Taq* DNA polymerase using the plasmid, primer, dNTP concentration as above, and 2.5 U EpiMark Hot Start *Taq* DNA polymerase in 1X EpiMark Hot Start *Taq* Reaction Buffer (and cycled 30 s at 95°C, 20 cycles of 15 s at 95°C, 15 s at 55°C, 30 s at 68°C, with a final incubation of 1 min at 68°C). The amplified inserts were purified and size selected using AMPure XP beads with a 1st bead selection of 0.55X volume of beads and a second bead selection of 0.35X sample volume. Concentration and purity of the final fragments were assessed by Agilent Bioanalyzer (DNA 1000).

Ligation assemblies were prepared with each fragment at a 5 nM final concentration in 1× T4 DNA ligase buffer with 2.5 μL of NEB Golden Gate Assembly Enzyme Mix in a volume of 50 μL. Reactions were incubated 1 h or 18 h at 37°C, then heat inactivated 10 min at 65°C. Alternatively, reactions were carried out with 30 cycles of 1 or 5 min at 37°C and 1 or 5 min at 16°C, followed by a 5 min 65°C heat inactivation step. Ligation products were blunted using the NEB Quick Blunting kit, adding 10 μL 10X Quick Blunting buffer, 10 μL 1 mM dNTPs, and 4 μL Quick Blunting Enzyme Mix (final volume 75 μL) incubating 1 h at 25°C, followed by cleanup using Monarch^®^ PCR & DNA Cleanup Kit columns. SMRT adapters were ligated by adding 2 μL SMRTbell blunt adapter (20 μM, Pacific Biosciences) and 25 μL Blunt/TA Ligase Master Mix in a final volume of 50 μL, incubating for 1 h at 25°C, then purification with Monarch^®^ PCR & DNA Cleanup Kit columns. The adapter-ligated library was treated with 1 μL PreCR Repair Mix in 1X Thermopol buffer, 1 mM each dNTP and 0.5 mM NAD+ in a 50 μL total volume, incubated 20 min at 37°C, then cleaned up with Monarch^®^ PCR & DNA Cleanup Kit columns. Finally, libraries were treated with 50 U Exonuclease III and 5 U Exonuclease VII in 1X Standard Taq Polymerase buffer in a 50 μL total volume, and incubated 1 h at 37°C. Final libraries were purified twice using a 1X volume of Ampure PB beads (Pacific Biosciences). Average fragment size was estimated by Agilent Bioanalyzer DNA 12000 assay (typically 1800-2400 bp) and total DNA concentration (typically 2 - 4 ng/μL) was determined. Libraries were prepared for sequencing according to the Pacific Biosciences Binding Calculator Version 2.3.1.1 and the DNA/Polymerase Binding Kit P6 v2 using the Magbead OCPW protocol and no DNA control complex. Libraries were sequenced on a Pacific Biosciences RSII, 1 SMRT cells per library, with a 6 h data collection time.

Consensus sequences were built for fragment assembly libraries with the Arrow algorithm using ccs program from SMRT Link software. Each consensus sequence represents a result of ligating multiple Golden Gate fragments into a single assembly such that the resulting consensus reads are comprised of long fragments separated by short regions corresponding to ligation junctions. Given the ten known 300nt fragments, their coordinates and mapping direction in each consensus read from assembly libraries were determined using BLAST software. This information was then used to tabulate the frequency of pairwise ligation events and overall composition of assemblies. A number of filtering steps were applied to ensure integrity of the derived data. Any 300nt fragment was required to map entirely from the first to the last nucleotide in the consensus read. Additionally, only two types of ligation junctions were expected to be seen in consensus reads: junctions of length 4 corresponding to overhang ligation during assembly reaction and junction of length 8 corresponding to blunt ligation during SMRT library preparation workflow. A 1nt variation in length was permitted for each junction type to account for possible errors in the sequencing reads. If any of above conditions were not met, the resulting consensus read was excluded from the analysis. When a blunt ligation junction was detected in the consensus read, the entire read was split apart at such junctions.

### Golden Gate cloning of 12 and 24 fragment *lac* cassettes

Golden Gate assembly reactions consisted of 75 ng pGGA destination plasmid, 75 ng of each precloned DNA fragment (See Supplementary Data, Tables S3 and S4 for fragment and junction identities), 500 U T4 DNA ligase, and 15 U BsaI-HFv2 (BsaI isoschizomer) in a final volume of 20 μl (12 fragment assemblies) or 25 μl (24 fragment assemblies) unless otherwise noted in the text. Reactions were kept cold with the use of a pre-chilled aluminum cold block before transfer to a T100 thermal cycler (Bio-Rad). Assembly reactions were incubated with thirty cycles consisting of 5 min at 37°C and 5 min at 16°C, followed by a 5 min final incubation step at 55°C then a final 4°C hold prior to transformation. Transformations were performed using 2 μl of each assembly reaction added to 50 μl competent T7 Express cells, incubation on ice for 30 min, incubation at 42°C for 10 s, with a final 5 min recovery period on ice. SOC outgrowth medium (950 μl) was added and the cells were incubated 1 h at 37°C with rotation. The outgrowth was then placed on ice for 5 min before plating 5 μl (12 fragment assemblies) or 100 μl (24 fragment assemblies) using bead spreading on prewarmed agar plates (Luria-Bertani broth supplemented with 1mg/mL dextrose, 1 mg/mL MgCl2, 30 μg/mL Chloramphenicol, 200 μM IPTG and 80 μg/mL X-gal). Plates were inverted and placed at 37°C for 18 h, then stored at 4°C for 8 h before scoring colony color phenotype. Plates were imaged and counted using the aCOLyte 3 automated colony counting system (Synbiosis) or by hand. For each assembly type, total transformants and percentage correct assemblies (blue colonies) are reported as the average result of at least three independent assembly reaction replicates, with the reported error one standard deviation from the mean.

## RESULTS

In the current study, we have extended our previously described method for rapidly profiling DNA ligase end-joining fidelity and bias to substrate pools with randomized four-base overhangs. Libraries were prepared with ligation temperatures of 25°C or 37°C with ligation times of 1 or 18 hours. The multiplexed ligation profile results for overnight ligation (18 h) at 25°C are shown in Figure 2. Data visualization from other tested ligation conditions can be found in the Supplementary Data, as can the raw results tables for all conditions and analysis of reproducibility between replicates (Supplementary Data, Figures S1-S6 and Supporting Data). Four-base overhang ligation libraries showed increased yields with prolonged incubation, with 56±6% ligation at 1 h, increasing to 82±3 % yield at 18 h. However, fidelity for each overhang changed little from short to long incubation time, despite the increased library yield (Supporting Figure S1), indicating that mismatch ligation was occurring proportionately throughout the ligation time course. Overall fidelity was dramatically improved for four-base overhangs at 37°C (Supporting Figure S2 and S3), however, the bias in efficiency between overhangs was much more pronounced at 37°C than at 25°C, with high AT overhangs notably underrepresented compared to high GC overhangs. This sequence-dependent yield bias was reduced after 18 h incubation. The increased bias is reflected in lower ligation yields at this temperature, 45±2% product at 1 h increasing to 69±3 % at 18 h.

In discussing the ligation profile results, individual overhangs are written in the 5′ to 3′ direction with the phosphate omitted, and ligation products are written as overhang pairs with the top overhang written in the 5′ to 3′ direction and the bottom overhang in the 3′ to 5′ direction. For example, 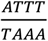 represents the fully Watson-Crick paired ligation product between a substrate with a 5′-pATTT overhang and a substrate with a 5′-pAAAT overhang.

Figure 2, Panel A shows a log-scale frequency heat map of all ligation events, with the overhangs sorted such that the top left to bottom right diagonal represents Watson-Crick paired ligation products. Panel B shows the linear frequency of ligation events for each overhang as a bar plot, with the Watson-Crick ligation frequency shown in blue, and the summed frequency of mismatch products shown in orange. While the majority of Watson-Crick paired ligation partners were observed in similar overall frequency (Figure 2B, blue bars), several overhangs had notably reduced incidence. The majority of the low-abundance overhangs were TNNA, with the corresponding ANNT overhangs not underrepresented. Several other high AT% overhangs, such as AAAA and TTTT, were also seen in reduced numbers as compared to the average overhang. This result mirrored our previous observation in three-base overhangs that TNA overhangs ligate significantly slower than the reciprocal ANT overhangs (21).

**Figure 2.**
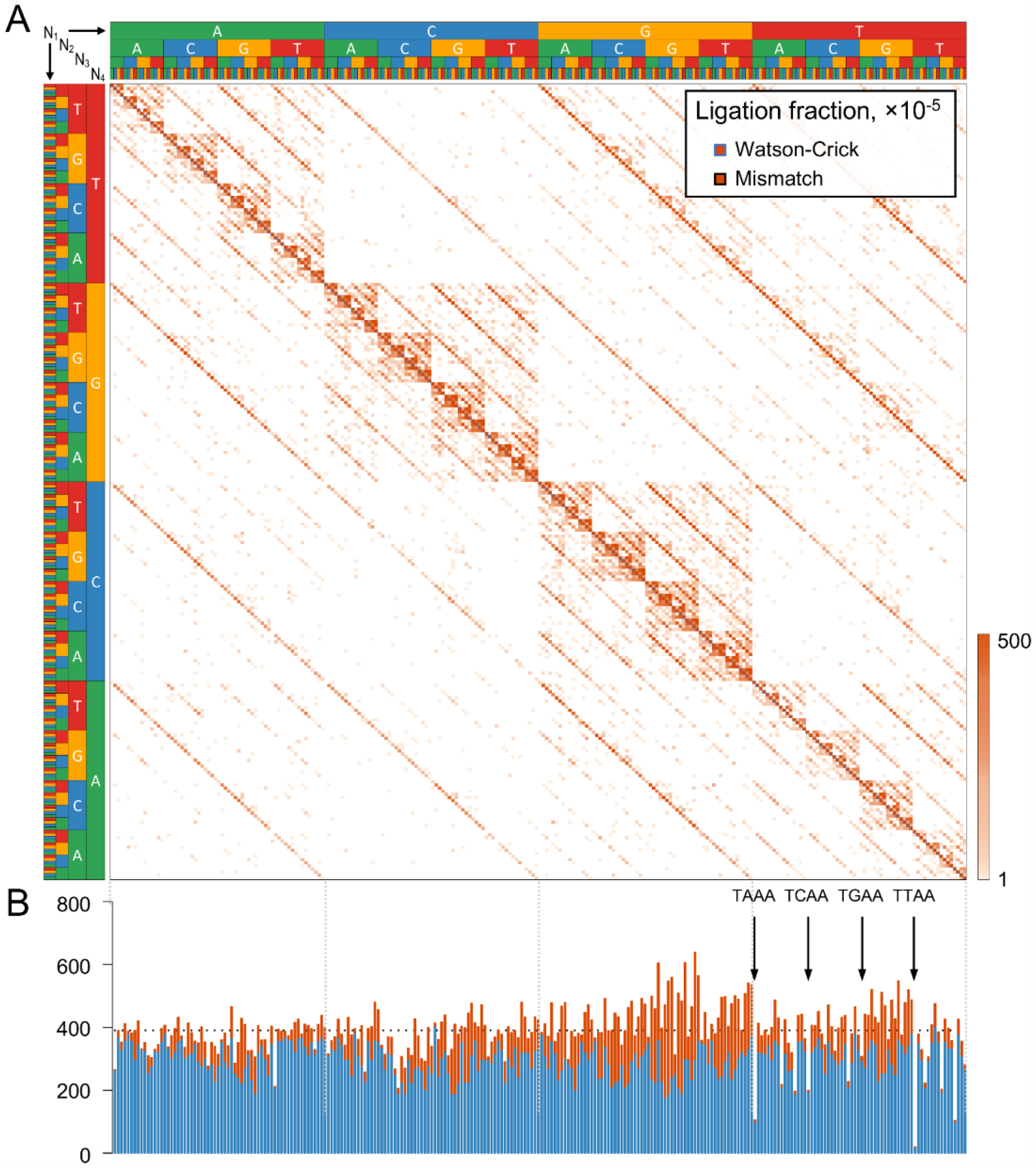
Assay results for the ligation of randomized four-base overhangs by T4 DNA Ligase. SMRT sequencing results for ligating 100 nM of the multiplexed four-base overhang substrate 18h at 25°C, with 1.75 μM T4 DNA ligase in standard ligation buffer. Observations have been normalized to 100,000 ligation events (see Supporting Data for actual observation totals). (A) Frequency heat map of all ligation events (log-scaled). Overhangs are listed alphabetically left to right (AAAA, AAAC…TTTG,, TTTT) and bottom to top such that the Watson-Crick pairings are shown on the diagonal. (B) Stacked bar plot showing the frequency of ligation products containing each overhang, corresponding to each row in the heat map in (A). Fully Watson-Crick paired ligation results are indicated in blue, and ligation products containing one or more mismatches are in orange.

The range of observed ligation fidelity as a function of overhang identity is quite broad, from overhangs with very few mismatch ligation events (e.g., AAGA, CCAA), to those where the majority of ligation partners had at least one base-pair mismatch (e.g., GGCG and GGCC). Overall, there was a weak trend towards lower fidelity with higher GC content. Additionally, 5′-GNNN and 5′-GGNN sequences were particularly overrepresented in the low-fidelity region. The position dependence of the specific mismatched base-pairings observed for the 25°C ligation temperature are visualized in Figure 3. For both the edge (N1:N4’, Figure 3A) and middle (N2:N3′, Figure 3B) positions, G:T mismatches were the most frequently observed ligation event, with a 5′G and a templating T being significantly more prevalent than a 5′T with a templating G (though far less dramatically than in the previously reported 3-base overhangs (21)). T:T mismatches were also common, as well as purine:purine mispairs. In the latter case, G:G mismatches dominated in the middle positions, while A:G and G:A mismatches were preferred at the edge position. At 37°C, the specific mismatches observed matched those seen at 25°C ligation, but were found in overall reduced incidence (Supplementary Data, Figure S7). For most overhangs, it should be noted that the bulk of the mismatch ligation events were derived from pairing a few specific ligation partners. This result suggested the possibility of even “low-fidelity” overhangs being used in high-fidelity ligation sets, as long as their most favorable mismatch ligation partners were not present.

**Figure 3.**
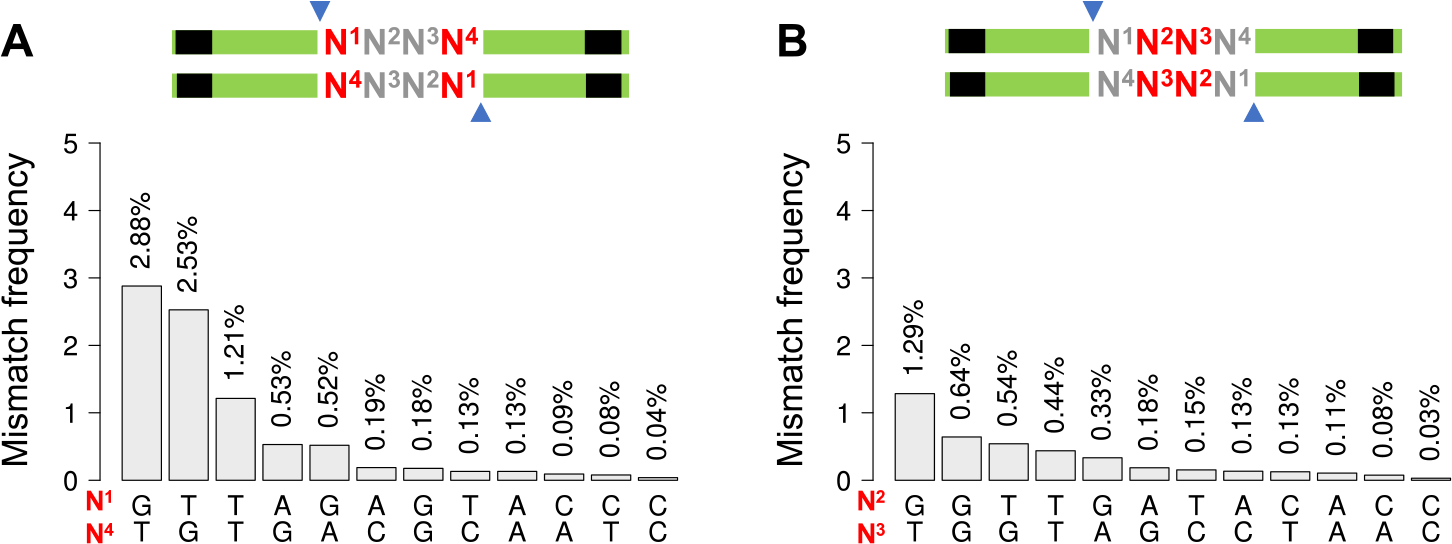
Frequency of specific base pair mismatches by position. Incidence of each possible mismatched base pair observed during ligation of four-base overhangs, with 100 nM of the multiplexed substrate, 1.75 μM T4 DNA ligase, and 18 h incubation at 25°C in standard ligation buffer. This figure was generated from the same data as shown in Figures 2. (A) shows the results for the edge position (N1:N4’); (B) for the middle position (N2:N3′).

### Ligation fidelity library is predictive of fidelity in multi-fragment gene synthesis

A key potential application of the ligation fidelity and bias data generated in this study was to predict the results of one-pot, multi-fragment assembly reactions. The fidelity profiles allowed the selection of sets of junctions that were predicted to be highly “orthogonal,” that is, having low potential for mismatch ligation events with any other member of the set. Thus, to test the predictive power of our ligation fidelity data, 10-fragment assemblies were designed for Golden Gate assembly protocols, picking junctions to give a variety of predicted assembly outcomes. Fragment inserts (A-J) consisted of 10 randomized 300 bp sequences flanked by specific four-base overhangs and BsaI restriction sites (Figure 4). For all assemblies, Insert A and J were terminated with an overhang not predicted to pair or mispair with any other overhang present in the assembly reaction (AAAC). The junctions between fragment pairs (Junctions 1 – 9) were selected to either be 9 high-fidelity (HF) junctions, or a low-fidelity (LF) set where 9 junction pairs were chosen such that many mismatch ligation events were predicted (Table 1). Two additional sets were designed: a deletion-prone (DP) set, where junction 7 of the HF set was changed to 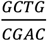 such that deletion (and to a lesser extent, duplication) of insert G was predicted to result; a failure-prone (FP) set where junction 7 was replaced with the high fidelity but low efficiency 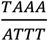 pair (Table 1).

**Figure 4.**
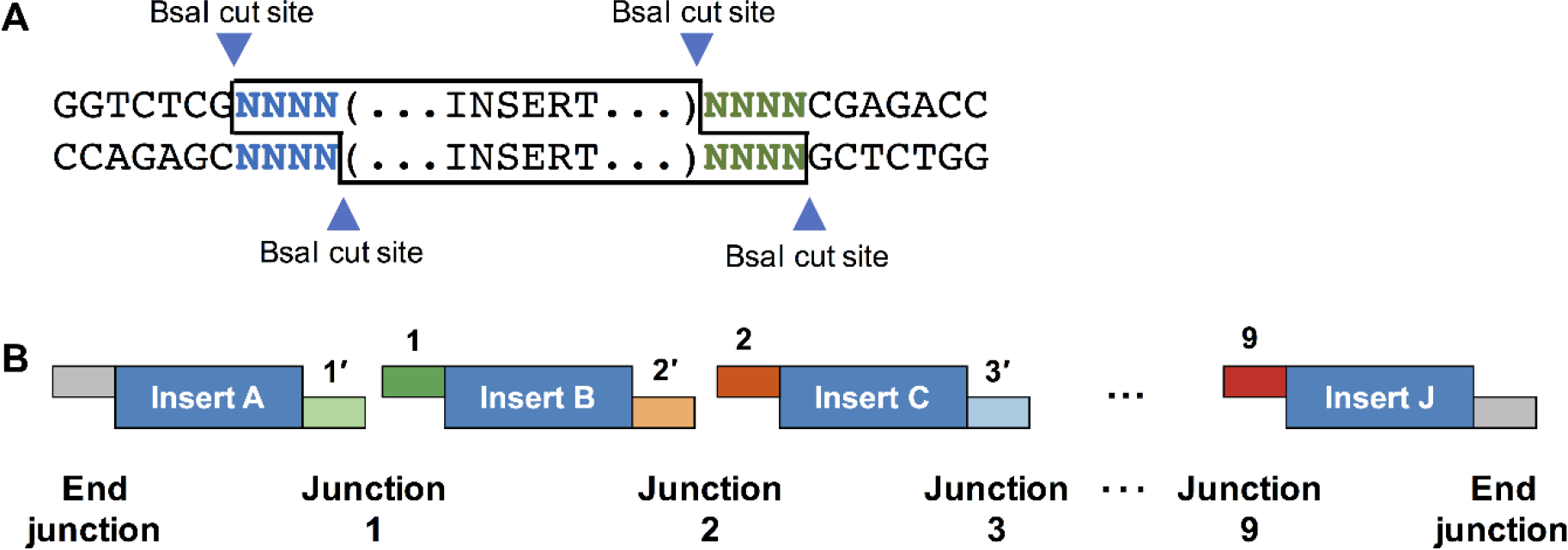
Overview of Golden Gate assembly design. Ten fragments of arbitrary, randomized sequence (Supplementary Information, Table S2) were designed, giving 9 junctions and an “end junction” designed with sequence AAAC, which was not predicted to have significant mismatch ligation potential with any overhang used for the junctions. The sequences chosen for the junction differ amongst the HF, LF, DP and FP sets, as indicated in Table 1. The order of assembly could be determined by SMRT sequencing of the products, with the unique insert sequences defining the order of assembly and thus, which overhangs ligated to produce the connection.

For each set of inserts, Golden Gate assembly was tested using typical cycling conditions (5 min 37°C, 5 min 16°C, 30 cycles), and the product assemblies (all lengths) were sequenced by SMRT sequencing. This method allowed identification of the number, identity, and orientation of fragments in each assembly (Figures 5, 6 and Supporting Data). The experimental results were compared to predictions based on the four-base overhang fidelity data collected under the same conditions. The HF set indeed assembled with high fidelity and high efficiency: 99.9% of all observed assemblies were formed by correct Watson-Crick pairings. Among assemblies starting with Insert A and ending Insert J (effectively, those that would be predicted to assemble into the destination plasmid), 99.9% contained all 10 fragments in the proper order and orientation under both reaction conditions. Shorter, incomplete assemblies were equally high fidelity, with most short fragments representing incompatible partial assemblies (e.g., ABCDEFG and GHIJ; see Supporting Data for a list of all fragments). For the DP set, we observed an increase in 9- and 11-fragment assemblies with an incorrect junction; analysis of the specific structures showed that these were dominated by deletion or duplication of fragment G, as predicted. The LF set showed dramatically increased numbers of erroneous junctions, with 74.1% of assemblies containing at least one erroneous junction. The FP set displayed a drop in ligation events at junction 7 (~30% decrease relative to other junctions, and there were many fragments observed truncated at this junction (ABCDEFG and HIJ). However, while this connection did appear to fail to form with increased frequency, it was not an impediment to seeing full length assemblies. Additionally, the HF, DP and FP sets all showed an increased incidence of truncations at junction 6 (the 100% GC junction GCCG/CGGC) and a drop of roughly 23% of connections at junction 6 relative to the average for all junctions, despite this sequence not being a predicted low efficiency junction in the ligation fidelity data set.

In addition to successfully predicting overall fidelity, the specific observed junctions were well predicted by the ligation fidelity library data (Figure 6). Low-frequency mismatch ligation events were not well predicted in any set (less than 10 observations per 100,000 ligation events). Observed erroneous junctions in the HF set seemed random, with some matching predicted mistakes and others not seen in the fidelity libraries at all. For the higher frequency predictions in the LF set, the expected junctions were observed, and largely in the frequency predicted by the fidelity data. Some junctions, however, were observed in greater or lower relative incidence than were predicted, and overall a moderately higher incidence of many mismatches than the fidelity library data gathered at 25°C would predict. Nevertheless, it was possible to qualitatively predict not only whether sets of nine junctions would be high fidelity or low fidelity in their assembly, but which specific mismatch ligation events would appear in the low-fidelity sets; in no case did we see significant incidence of misligations not predicted by the fidelity data, nor fail to see a mismatch that was predicted. The same was true of the predictions of the DP and FP sets (Supporting Figure S9).

Assembly reactions with prolonged incubation (18 h) at 37°C in lieu of cycling were also tested. The results (Supporting Figure S10) are largely consistent with the results of cycling with a few key exceptions. Firstly, the incidence of ligation errors in the LF set was much lower with the higher ligation temperature, consistent with the observation that fidelity was much improved at 37°C in the multiplexed fidelity profiles. While the HF set assembled under these conditions again showed >99.9% of all observed assemblies formed by correct Watson-Crick pairings, the LF set had only 31.2% of all assemblies containing at least one mispair; the specific mismatches observed were the same, just present in lower prevalence. Overall, the cycled assembly conditions were in good agreement with predictions made by the fidelity library ligated at 37°C. This result indicates that if a suitable orthogonal set of overhangs is chosen, either method will result in high fidelity assembly, but if sets prone to mismatch ligation events are chosen, a ligation temperature of 16°C will result in significantly more failed assemblies. Under the high temperature incubation conditions, while the FP still showed a large increase in truncations at the predicted low-efficiency junction 7, the increase in truncations at the 100% GC junction 6 was much less noticeable.

### Use of fidelity predictions enables 12- and 24-fragment one pot assembly of *lac* cassettes

We next sought to test the predictive power of our ligation fidelity data in a practical application to select DNA fragment breakpoints and overhang sequences. Thus, we designed 12- and 24-fragment Golden Gate assemblies of a cassette containing both the *lacI* and *lacZ* genes, as well as necessary regulatory elements to drive expression of β-galactosidase (β-gal); see Supplementary Data for full cassette sequence, the coordinates of the break points chosen for all fragments, and sequences of holding plasmids and the pGGA destination vector containing the chloramphenicol resistance gene (Supporting Data and Tables S3 and S4). Easily scored blue colonies indicate fully assembled sequences and an intact open reading frame for *lacZ*. White colonies indicated an intact pGGA plasmid containing the chloramphenicol resistance gene, but with the *lac* cassette not faithfully assembled and thus, no β-gal expression. Each of the fragments comprising the *lac* cassette were approximately equal in length, and were pre-cloned into vectors flanked by BsaI restriction enzyme recognition sequences. The fidelity data was applied to select junctions for 12- and 24-fragment test systems that were predicted to be orthogonal sets that should assemble with high fidelity, without modification of native sequence. Additionally, we designed a deletion-prone 12-fragment test system with overhangs predicted to frequently result in a deletion within the *lac* cassette, but still result in assembly of many circular constructs granting chloramphenicol resistance. The predicted fraction of correctly assembled lac cassette-containing circular plasmids to those which would circularize but be missing one or more fragments was 99% and 91% for the high fidelity 12- and 24-fragment sets, respectively, and 33% for the low fidelity 12-fragment set.

Assembly of the high-fidelity 12-fragment test system using the typical cycling protocol resulted in large numbers of transformants, with ~99% harboring plasmids with accurate assemblies based on blue/white scoring (Figure 7A and Supplementary Data, Table S6), closely agreeing with predictions. Exclusion of any of the 12 fragments resulted in plates with no observed blue colonies (Supplementary Data, Table S7). Use of the predicted low-fidelity, deletion-prone test system resulted in 45% ± 5% of the transformants harboring correct assemblies, in good agreement with the predicted frequency of ~33% (Figure 7B, Supplementary Data, Table S6). The assembly of the predicted high-fidelity 24-fragment test system resulted in a lower count of transformants (~1 transformant/μL of assembly mix, vs ~100/μL for the twelve-fragment assembly), likely due to the increased number of junctions. However, the observed frequency of transformants expressing β-gal was still quite high, 84 ± 5%, only modestly lower than the predicted 91% of correct assemblies based on the fidelity data. As in the 12-fragment assembly, omitting any one of these fragments eliminated the blue phenotype on scored plates (Supplementary Data, Table S8). Thus, use of the fidelity data to guide choice of junctions leads to the ability to flexibly use native sequence while still subdividing the assembly into >20 fragments that can be assembled accurately in a single assembly reaction.

### DISCUSSION

Herein, we report the comprehensive fidelity and bias profile of T4 DNA ligase in the joining of four-base overhangs, and have applied the results to the accurate prediction of high-fidelity fragment sets for Golden Gate assembly. The fidelity of four-base overhang ligation shares much in common with our previously reported data for three-base overhangs. As in the case of three-base overhangs, G:T mismatches were highly favored, along with lesser amounts of T:T, purine:purine, and A:C mismatches. However, here, very similar profiles were seen at the edge and middle positions, in contrast to three-base overhangs in which mismatches were dramatically dis-favored at the middle position, and only T:T mismatches were observed. Four-base overhangs also lack the dramatic asymmetry in preference for 5′-purines; while G:T is favored modestly over T:G (T in the template vs G in the template), it falls far short of the 10-fold preference observed for three-base overhangs. Thus, the data is suggestive of a stronger influence of annealing on the mismatch preferences as compared to three-base overhangs. However, the influence of ligase preference is still clear, with higher mismatch frequencies at the edge position and sequence dependence of mismatch prevalence. For example, cases of an edge mismatch with two GC and one AT base pair where both GCs are in the middle (N2:N3′ and N3:N2’) positions outnumber by a factor of two those where only one of the middle pairs is a GC.

While the trends in the ligation fidelity profile data presented here are informative as to the overall mismatch tolerance of T4 DNA ligase, practical utility can also be extracted from the knowledge of precisely which mismatched four-base overhang pairs will ligate and which will not. This data set can be used to enumerate sets of “orthogonal” Watson-Crick pairs of overhangs, predicted to have little to no side products from mismatch ligation events. This application allows for empirical choice of overhangs for use in Golden Gate type assemblies that goes beyond the rules of thumb. In short, it is not necessary to exclude all pairs that have only a single mismatch, as many of these (e.g. 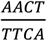,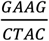) form very few if any mismatch ligation products with each other or with their complements. The data sets can be used to select a very large number of sets of 10, 12, or even 20+ Watson-Crick pairs with low levels of cross-talk. Further, the fidelity data allows identification of low-efficiency Watson-Crick pairs, allowing expected inefficient sets to be excluded from assembly design.

However, it should be noted that while assemblies used in this study matched predictions very closely, Golden Gate assembly requires cutting by a Type IIS restriction enzyme and melting of the overhangs, steps that are not directly studied in the multiplexed ligation profiling assay. Thus, overhangs that lead to slow melting, i.e., 100% G/C overhangs, may not assemble as efficiently as predicted from ligation data alone. Indeed, in the observed data from the thermocycled assembly conditions (5 min 16°C, 5 min 37°C, 30 cycles, Figure 5 and Supporting Data) the frequency of ligation events observed at junction 6 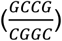 lagged behind predictions, while other junctions lined up much better with predicted efficiencies. Under the assembly conditions of 37°C 18 h (Supplementary Data, Figure S8), the 100% G/C pair did not result in an undue number of assemblies that were truncated at junction 6. Thus, when lower ligation temperatures are used, the reduced efficiency of high GC overhangs may remain an issue in assembly reactions not fully accounted for by the predictions. Under both conditions temperatures, predicted low efficiency (especially TNNA) junctions need to be excluded to avoid a reduced yield due to the presence of truncation products.

**Figure 5.**
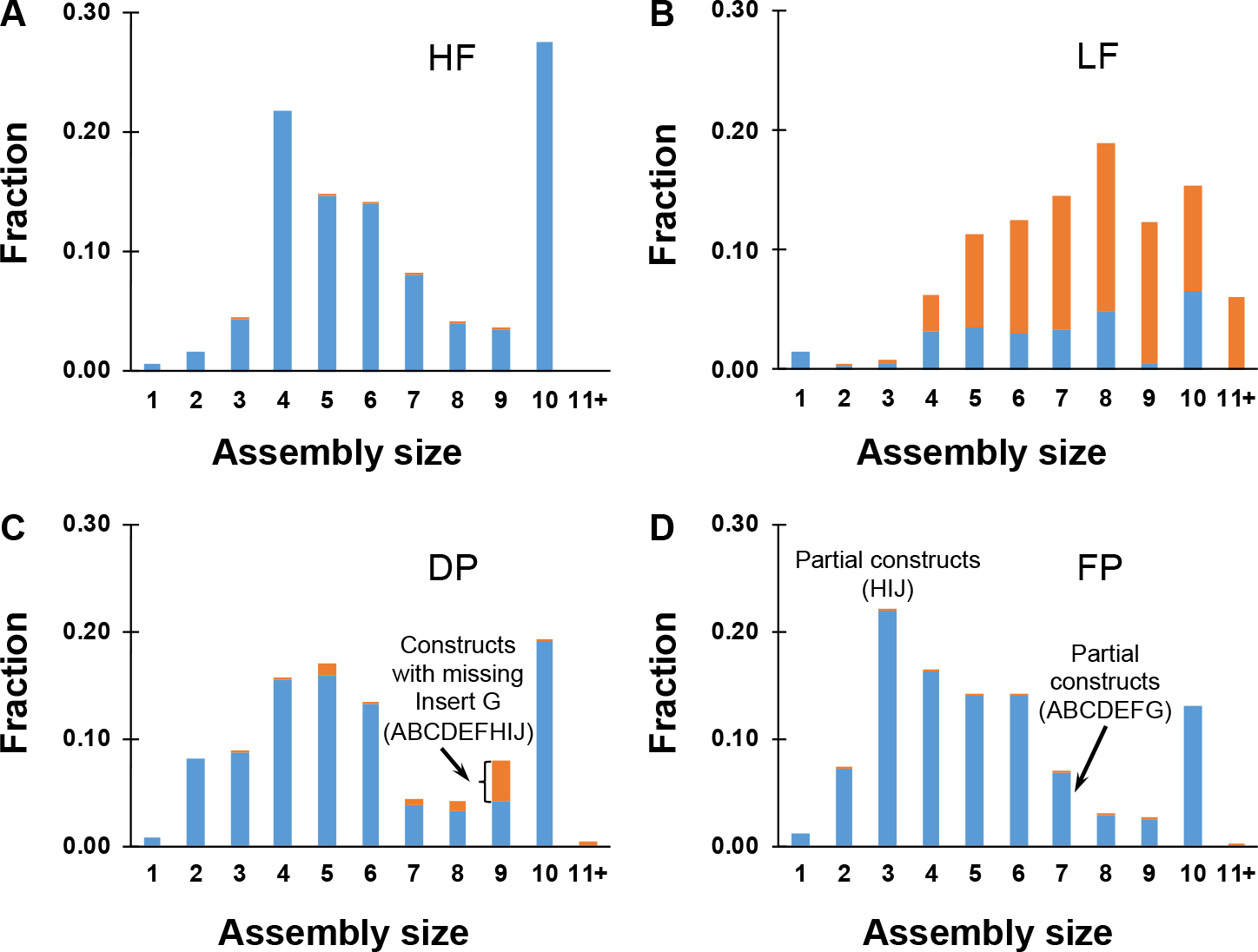
Distribution of assembly sizes for the 10-fragment Golden Gate assemblies (37°C 5 min/16°C 5 min, 30 cycles). Correct constructs are in blue, constructs containing at least one incorrect junction are shown in orange. (A) HF set results in correctly assembled constructs with the full-length product ABCDEFGHIJ being the most common. (B) LF set results in a significant fraction of incorrectly assembled constructs as expected. (C) DP set leads to accumulation of a construct with missing Insert G and a slight uptick in 11-insert assemblies duplicating fragment G. (D) FP set has a ligating junction 7 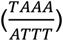 with predicted low efficiency; this junction did join, but with ~33% reduced incidence as compared to the other junctions. Additionally, many product fragments truncated at this junction (ABCDEFG and HIJ) were observed (Supporting Data).

**Figure 6.**
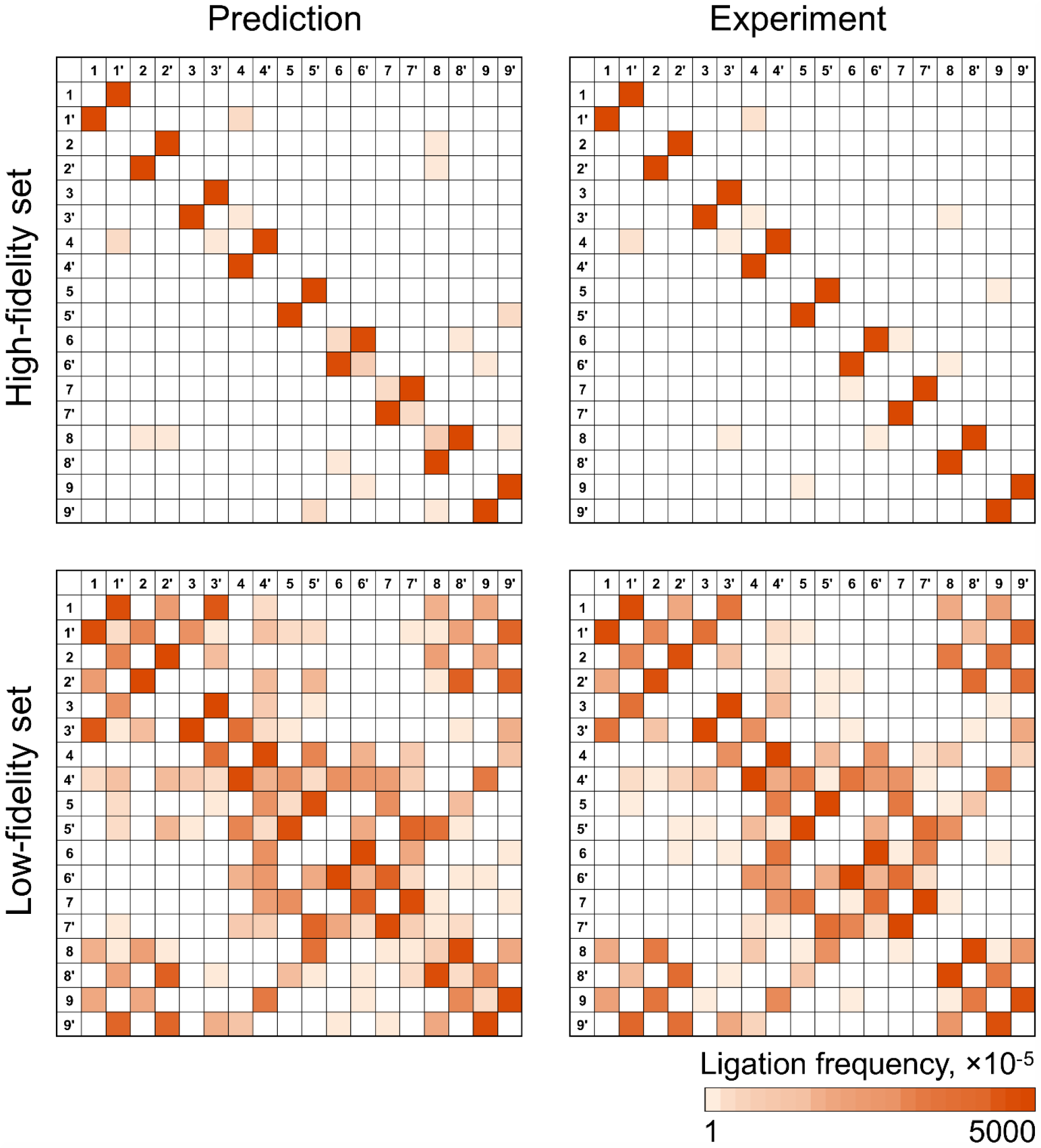
Predicted versus observed fragment linkages in Golden-Gate assembly of the HF and LF 10-fragment assemblies. Junction overhangs can be found in Table 1. The intensity of the color corresponds to the number of instances of that junction observed in a Pacific Biosciences SMRT sequencing experiment, normalized to 100,000 total junctions. Predicted frequencies of junctions are based on the fidelity library data generated for the four-base overhang substrate ligated with T4 DNA ligase at 25°C for 18 h. The experimental observations shown are for assembly of the 10-fragment HF and LF sets with Golden Gate Assembly mix, 37°C 5 min/16°C 5 min, 30 cycles.

**Figure 7.**
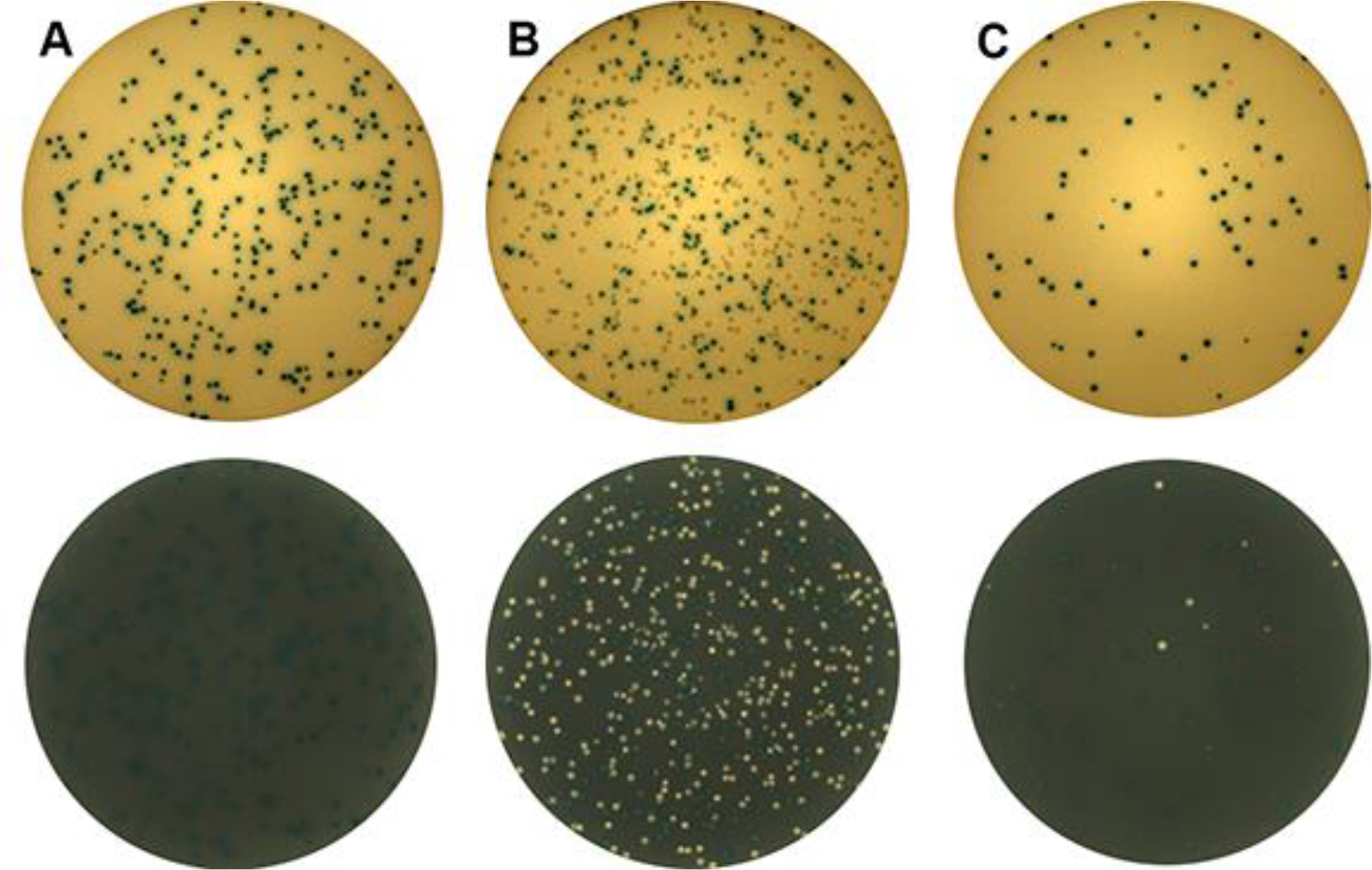
Twelve- and 24-fragment Golden Gate assembly of *lac* cassettes. Assemblies were twelve (A and B) or twenty-four fragments (C) in a single pot, with choice of junctions driven by the ligation fidelity and bias profile. Reactions were performed as described in the Materials and Methods section, plating 5 μL assembly reaction for 12-fragment assemblies, and 100 μL for 24-fragment assemblies. Plates shown are representative replicates, imaged and counted using the aCOLyte 3 automated colony counting system with a white filter (top) to show blue colonies expressing β-gal, and a black filter (bottom) to visualize white colonies containing antibiotic resistance but a non-functional *lac* cassette. Data for all replicates can be found in the Supplementary Data, Table S6. (A) shows the results of a designed predicted high fidelity 12-fragment set, predicted 99% blue colonies, observed average over 8 replicates, 99.2 ± 0.6 %. (B) shows results of the designed low fidelity, deletion-prone 12-fragment set; predicted 31% blue colonies, observed average of 8 replicates, 45 ± 5%. (C) shows the results of assembly of the designed 24-fragment high fidelity set, predicted 91% blue colonies, observed average over 10 replicates, 84 ± 5%.

The data set that best predicted the results for the cycled assembly conditions (5 min 16°C, 5 min 37°C, 30 cycles) was the multiplexed ligation profile data collected at 25°C, 18 h incubation time. From this data, we went on to see if the predictions would hold for actual gene assemblies, requiring circularization and successful transformation. To this end, the 12- and 24-fragment lac cassettes were designed and assembled. The results matched predictions remarkably well, showing that native coding sequences can be divided into at least twelve fragments and still assemble with effectively perfect accuracy in one pot. The 24-fragment assembly showed more errors, expected based on predictions, but still achieved a remarkable 80% of transformants expressing β-gal. This result suggests that even higher accuracy could be achieved by removing incomplete assemblies before transformation. However, the current results indicate that assemblies of more than 20 fragments in one pot can be achieved with high accuracy.

When junctions can be arbitrarily chosen, very large assembly sets with high fidelity should be possible. For use in cases where junctions are not restricted by coding sequence, we have enumerated several sets of overhangs predicted to have negligible mismatch-ligation cross talk (>98% correct ligation events). Set 1 (Table 2) contains 15 overhang pairs that includes the MoClo standard overhangs (13,14); this set is preferred when at least some of the parts have already been designed with this standard in mind. If users are not restricted by MoClo, set 2 provides a set of 20 overhang pairs predicted to achieve the same overall fidelity. To ensure efficient assembly, these two sets exclude all Watson-Crick pairs that ligates significantly below the mean. We have additionally excluded any overhangs with 100% GC content, which should remove concerns with inefficient melting of these sequences under the cycled protocol. Finally, set 3 enumerates 25 Watson-Crick pairs based on the ligation profile at 37°C 18h; at this temperature assembly is higher fidelity overall, allowing many more overhangs to be used while still predicting >98% assembly fidelity. Fully GC overhangs do not appear to be a concern under these conditions, but more overhangs were excluded due to low ligation efficiency in the ligation profile data. The Sets 1 and 2 are best used for cycled 16°C/37°C assemblies; Set 3 for prolonged (overnight to ensure high efficiency ligation) static incubation at 37°C.

**Table 2.** Predicted high fidelity four-base overhang sets for use with Golden Gate assembly methods. Predicted high fidelity four-base overhang sets for use with Golden Gate assembly methods. Sets are provided for use with cycled assembly (16°C/37°C cycles; Sets 1, 2) and a set for use with static incubation at 37°C (Set 3). Set 1 is an extended MoClo set (TGCC, GCAA, ACTA, TTAC, CAGA, TGTG, GAGC) with additional 8 overhangs. All sets are predicted to assemble with a >98% overall fidelity if every overhang and its complement is used; subsets of these sets are predicted to have even higher fidelity.

| **Set** | **Number of**<br>**overhangs** | **Overhang sequences <sup>1</sup>** |
| --- | --- | --- |
| 1 | 15 | TGCC, GCAA, ACTA, TTAC, CAGA, TGTG, GAGC, AGGA, ATTC, CGAA, ATAG, AAGG, AACT, AAAA, ACCG |
| 2 | 20 | AGTG, CAGG, ACTC, AAAA, AGAC, CGAA, ATAG, AACC, TACA, TAGA, ATGC, GATA, CTCC, GTAA, CTGA, ACAA, AGGA, ATTA, ACCG, GCGA |
| 3 | 25 | GCCC, CCAA, ATCC, GGTA, ACGG, AAAT, ATAG, CTTA, AGGA, AGTC, ACAC, ATGA, GCGA, CATA, CTGC, AACG, CGCC, AGTG, CCTC, GAAA, CAGA, ACCA, AAGT, CGAA, CAAC |

<sup>1</sup> Only one sequence for each overhang pair is shown.

Any subsets of these sets are also predicted to be very high fidelity with matched efficiency, giving a great deal of flexibility to junctions that can be used in a given Type IIS-based assembly reaction. Additional high fidelity sets can be enumerated from the data (provided in full in Supplementary Data), and can be used to guide division of any large DNA construct to be divided into an arbitrary number of approximately equal fragments by choosing junction points predicted to form a highly orthogonal set. As an additional possibility, the ability to accurately predict specific mismatch-prone junctions (as in the DP set) could allow design of Golden Gate assembly sets that allow for *in vitro* “alternative splicing” of constructs. For example, as in the DP set, a 10-fragment assembly could be designed such that a certain fraction of assemblies will be missing one or more fragments based on the use of a deletion-prone overhang pairing. As a final note on joining accuracy, while a significant concern for Type IIS restriction enzyme-dependent methods, overhang mispairing is not predicted to be an issue in traditional restriction-enzyme based cloning methods utilizing Type IIP palindromic cutters. Only trace cross-talk is predicted between all possible palindromic overhangs (Supporting Figure S11), though our data suggests experimenters should avoid the poor-reacting TTAA and TATA overhangs.

The current method has proven effective in rapidly profiling the ligation fidelity of T4 DNA ligase in a single experiment. The data generated has allowed us to accurately predict the efficiency and fidelity of assembly reactions of up to 24 fragments. Further application of the method will allow for profiling the effect of other ligases, buffers, and protocols on ligation fidelity and bias. These data will allow for discovery of high fidelity, low bias ligation conditions that could extend the utility of Type IIS restriction based assembly systems even further. Finally, modifications of the substrate to include the restriction cleavage and melting steps should increase the accuracy of predictions and allow co-screening of different Type IIS restriction enzymes and ligases in combination. Thus, by combining informatics to guide junction choice and high-throughput screening of conditions, the use of dozens of fragments in a single pot, resulting in highly efficient and highly accurate assembly, is within reach.

## AVAILABILITY

Sequencing data pertaining to this study has been deposited into the Sequencing Read Archive under accession numbers SRP144368 (multiplexed ligase fidelity sequencing data) and SRP144386 (golden gate sequencing data). Custom software tools are available in the GitHub repository at: https://github.com/potapovneb/golden-gate.

## SUPPLEMENTARY DATA

Supplementary Figures S1-S11, Supplementary Tables S1-S8, Excel and .csv formatted data tables for raw ligation product observation counts, and 10-fragment Golden Gate assembly reactions. Supplementary Data are available online as a separate file.

## ACKNOWLEDGEMENT

We would like to thank Laurence Ettwiller, Laurie Mazzola, Rick Morgan, Yvette Luyten (NEB) and Pacific Biosciences for assistance with sequencing reactions. We are grateful to Bill Jack, Andy Gardner, and Karen Lohman for critical feedback on this manuscript.

